# Conflicting binocular input triggers inhibition followed by rebound, explaining paradoxically fast reaction times

**DOI:** 10.64898/2026.03.31.715537

**Authors:** Gábor Horváth, János Radó, András Czigler, Diána Fülöp, Sári Zoltan, Ilona Kovács, Péter Buzás, Gábor Jandó

## Abstract

Binocular vision depends on the integration of matching visual features across the two eyes, while conflicting interocular signals can engage active inhibitory processes in the visual system. To investigate the temporal dynamics of these putative inhibitory processes, we examined how transitions between different binocular correlation states influence perceptual detectability and response speed. Using dynamic random-dot correlograms - free of monocular cues and allowing precise interocular manipulation - we presented brief target intervals embedded in longer background sequences. Stimuli varied in binocular correlation: correlated (C) patterns contained identical luminance profiles in both eyes, anticorrelated (A) patterns had inverted luminance dots, and uncorrelated (U) patterns had independent dot arrangements.

Across three experiments, we measured (1) the presentation duration threshold required to detect a change in correlation, (2) simple reaction times (RTs) to the same transitions at suprathreshold levels, and (3) psychometric functions across durations for selected transitions. In Experiment 1, A→C transitions yielded significantly higher duration thresholds than C→A, indicating a suppressive influence associated with prior anticorrelation. In contrast, Experiment 2 showed that A→C transitions produced the shortest RTs, while C→U transitions were slowest, suggesting a rebound-like facilitation following prior suppression. Experiment 3 confirmed these temporal and contrast dependences, with opposite changes in contrast threshold and reaction times between transitions toward and away from the correlated fusional states.

This divergence between perceptual onset and reaction time is consistent with a two-phase account in which binocular anticorrelation is associated with an initial suppressive phase followed by rebound-like facilitation that accelerates responses once the target becomes detectable. These findings are consistent with current models of binocular rivalry and fusion, and provide a temporally resolved behavioral perspective on how inhibitory control in sensory systems may dynamically influence subsequent responsiveness under conditions of perceptual ambiguity.

## Introduction

Stereopsis is one of the important visual faculties humans use to perceive the spatial layout of their environment. Horizontal disparity plays a crucial role in this process, arising from the slightly different viewing positions of the two eyes. To calculate disparity, the visual system must identify image points on the retinas that originate from the same locations on an object viewed from different angles - a challenge known as the correspondence problem (1,2). This problem essentially involves how the visual system matches these points to drive vergence eye movements and compute disparity. Throughout this process, the system identifies multiple potential matches - some correct and others false - which are later discarded in the visual processing hierarchy.

The initial solution relies on correlating the luminance profiles of the two images across various spatial frequency channels. If the profiles match, vergence movements are initiated to achieve fusion within the narrow depth range of Panum’s fusional area. Conversely, if fusion is unattainable - due to excessive differences between the images or disparities outside Panum’s area - rivalry or diplopia occurs (3).

While solving the correspondence problem is relatively straightforward in natural scenes - owing to the abundance of monocular cues - it becomes considerably more complex with Julesz’s random dot stereograms (RDS) (4). The ability to perceive depth in these stimuli, despite the absence of monocular information, demonstrated that disparity extraction can occur at early stages of visual processing. Julesz proposed - and subsequent physiological and psychophysical studies confirmed - that the neural substrates of these bottom-up processes reside in early visual cortex, preceding the integration of monocular cues such as edges, color, form, or motion (5–9).

Random-dot stimuli allow precise control over interocular similarity while eliminating alternative depth cues. This makes them particularly well suited for studying how the visual system handles conflicting binocular information. Neurophysiological studies have shown that neurons in primary visual cortex respond robustly to disparities in anticorrelated random dot patterns even though such stimuli do not normally support a stable depth percept, indicating a dissociation between early disparity encoding and later perceptual interpretation (1–3,10). These findings suggest that the brain explicitly represents both matched and mismatched interocular signals, and that additional mechanisms are required to resolve the resulting conflict at the perceptual level. While early visual areas encode anticorrelated disparities, higher visual areas show reduced or absent selectivity for such signals, suggesting the presence of later-stage mechanisms that suppress or reinterpret mismatched binocular input (4,5).

Psychophysical evidence also supports the idea of partially distinct binocular mechanisms, with interocular differences processed by specialized “differencing” channels rather than by the summation mechanisms that support fusion (6).

Low-level correlation mechanisms, also known as spatial cross-correlation, are optimally studied using random dot correlograms (RDCs) consisting of dots with binary luminance values. In these stimuli, the degree of correlation between the images - and thus the similarity of neural inputs to the two eyes - can be varied arbitrarily. If the luminance profiles are identical, there is a 100% luminance match and a correlation coefficient of +1 (correlated, ‘C’ state). When all corresponding dots have opposite luminance values, this results in a 0% luminance match and a correlation coefficient of −1 (anticorrelated, ‘A’ state). When two random dot images are generated independently, the correlation coefficient is 0 and only half of the dots share the same luminance (uncorrelated, ‘U’ state). Correlated images promote binocular fusion, whereas anticorrelated images typically evoke binocular rivalry or suppression.

Interocular correlation mechanisms have been probed by measuring the time needed to detect brief changes among correlation states in dynamic random-dot arrays. In a classic study, Julesz and Tyler presented three states (C, U, A) and measured the presentation duration necessary to detect brief changes between these states. Duration thresholds were significantly longer when the transition was from a non-correlated to a correlated state than in the reverse direction. These findings led Julesz and Tyler to introduce a phenomenological model involving two essential components: “neurontropy”, an entropy-like measure intended to characterize the degree of order in binocular neural activity, and a pair of antagonistic mechanisms that support fusion of matched signals and suppression or rivalry for mismatched signals, respectively. The latter has become a core ingredient of contemporary models of binocular vision emphasizing dynamic interactions between excitatory and inhibitory processes and competition between alternative interpretations of binocular input (12,13). Recent neurochemical imaging results further suggest that binocular conflict can alter the balance between excitation and inhibition in visual cortex, consistent with the involvement of active inhibitory control during the processing of mismatched binocular signals (14). The temporal dynamics of these interactions at the behavioral level, however, remain incompletely understood.

In a previous study, we detected variations in simple reaction time (RT) as a function of disparity and contrast (20). In the current paper, we ask whether transitions between different binocular correlation states reveal dissociable temporal signatures in perceptual detectability and reaction time. By combining duration threshold measurements with reaction time analysis within the same stimulus framework, we aim to characterize not only whether different correlation states interact, but also how their influence unfolds over time. Perceptual detection and response initiation are known to reflect partially distinct stages of sensory evidence accumulation and decision formation (15), which may be differentially influenced by preceding neural states.

We were particularly interested in further investigating the role of inhibitory interactions that may dominate between rivalry and fusion states. These questions are relevant beyond stereopsis research. Binocular anticorrelation provides a controlled model of interocular conflict, allowing investigation of how the brain resolves inconsistent sensory input over time. The observed dissociation between duration thresholds and reaction times suggests that inhibitory interactions do not merely suppress perception but dynamically shape subsequent processing. Such state-dependent modulation may reflect more general principles of sensory conflict resolution, excitation-inhibition balance, and perceptual decision-making in the brain.

## General methods

### Participants

The study was approved by the Regional and Institutional Research Ethics Committee of the Clinical Centre, University of Pécs (approval No. 5638) and was conducted in accordance with the World Medical Association Declaration of Helsinki (October 2013 revision). Participants were recruited during the period May 1, 2015 to December 31, 2020. All participants provided their written consent before completing the experiments. The observers had normal or corrected-to-normal vision, and random dot stereotests (16,17) confirmed appropriate binocular vision.

### Apparatus

A polarized flat LED 3D monitor (LG Cinema D2343P, refresh rate: 60Hz) with circularly polarizing goggles provided by the manufacturer was used for dichoptic stimulus presentation in all cases. The experiments were carried out in a darkened room. The head of the subject was placed on a chin rest at 100 cm viewing distance. The screen was masked on both sides, and the central region subtending 16°16° (1080×1080 pixels) served stimulus presentation. A fixation dot of 36 min of arc (’) diameter was presented in the center.

### Stimulus calibration

A high precision photometer (IL-1700 Photometer, International Light Technologies, Peabody, USA) was used for monitor calibration (18). The aims of the calibration procedure were: 1) to maintain constant mean luminance of 30 cd/m2 for all conditions, 2) to provide control over stimulus contrast and 3) to avoid monocular cues by minimizing interocular differences in luminance or contrast. As described earlier (18), this was achieved by adjusting the gray levels of the dark and bright dots independently for each channel using a custom made iterative least squares algorithm. For this procedure, a high precision photometer (IL-1700 Photometer, International Light Technologies, Peabody, USA) was used to measure the luminance characteristics of the left and right eye channels of the monitor.

The calculated RGB values were tested psychophysically by presenting concurrent vertical and horizontal square wave gratings in the left and right eyes, respectively. The gray levels of the dark and bright stripes were fine-tuned when necessary to ensure that only the intended grating was visible when viewed monocularly. The same criterion was employed to establish the optimal viewing position of the participant before each session.

### Stimuli

Dynamic random dot correlograms (DRDCs) served as stimuli that consisted of 50% bright and 50% dark square shaped 4×4-pixel dots that subtended 3.65’×3.65’. The random dot images were refreshed at 60Hz. The space averaged mean luminance was kept at around 30cd/m^2^. Three types of correlation were used to set up the stimuli: 1) correlated (C), 2) uncorrelated (U) and 3) anti-correlated (A). The correlation coefficients of the luminance profile between the left and right images were +1, 0 and -1 for C, U and A, respectively. Therefore, dots were identical in the two images for the C, their luminance was inverted for the A (i.e., no luminance match) and 50% were correlated and 50% were anti-correlated between the left and right images in case of the U type.

Each trial was composed of a background and a target image. In the case of the background, the stimulus area was completely filled up with one of the correlogram types (i.e., C, U or A). The target stimuli were patterns in which half of the area was filled with the background type of correlogram, while the other half was filled with another correlation state, called the target state. The pattern was either a checkerboard (Experiment 1, **Figure 1**) or stripes (Experiment 3). Importantly, the patterns could only be recognized by the observer if they perceived the transition from the background state to the target state binocularly. When viewed monocularly, subjects saw only random spatio-temporal noise for all background and target conditions.

**Figure 1.**
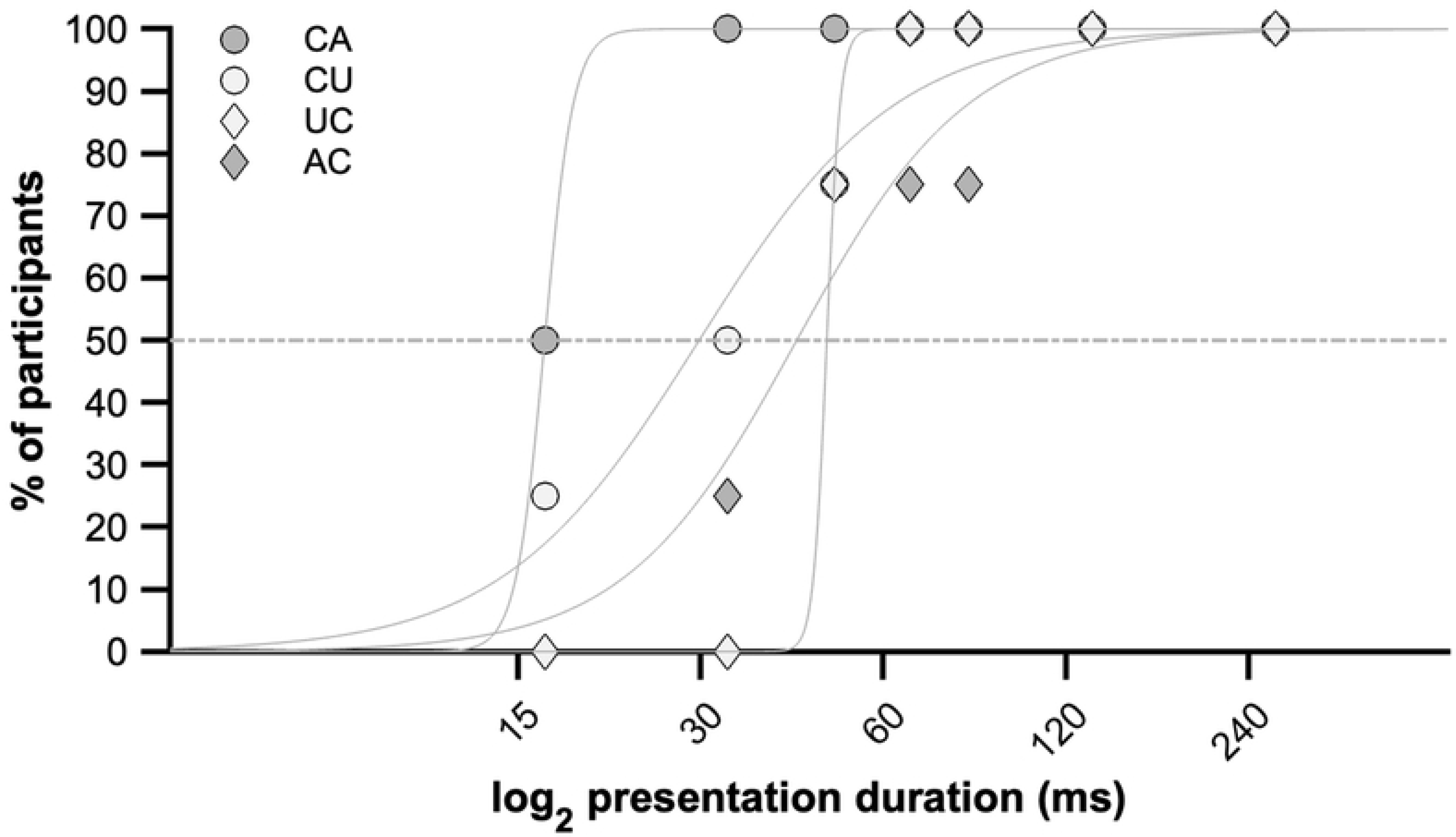
Types of binocular correlation changes used in the study. Experimental stimuli were composed of screen regions filled with dynamic random dot correlograms (DRDCs), whose binocular correlation was either correlated (C), or uncorrelated (U) or anti-correlated (A). These regions are illustrated by rectangles of different shading. The target stimulus was always a change in binocular correlation between a uniform background and a pattern composed of the background and a second correlation type. This resulted in 6 types of correlation changes (CU, UC, CA, AC, UA, AU). In Experiments 1 and 2, the target pattern was a checkerboard, whereas in Experiment 3, either horizontal or vertical stripes were used. The stimuli were presented on a 3D monitor and viewed through polarizing goggles. The target patterns could only be recognized when viewed binocularly and if the viewer had normal stereopsis.

Out of the six possible ones, four background-to-target transitions were tested in the present study: CA, AC, CU and UC. The letters indicate the correlation states, and their order specifies the direction of the transition. AU and UA transition conditions were omitted, because they were not detectable in the pilot measurements at 333ms presentation duration. The stimuli and tested conditions are demonstrated in **Figure 1**.

### Data analysis

SPSS 27.0.1.0 (SPSS Inc., Chicago, USA) was used for statistical analysis. Repeated measures ANOVA (rANOVA) was used to analyze the main effect of stimulus transitions and contrast. Reported are F-values, p-values and effect sizes (r). Mauchly’s test (test of the assumption of sphericity) and Levene’s test (test of homoscedasticity) were performed. Bonferroni-correction was applied for pairwise comparisons. For the analysis, p<0.05 significant level was set in all cases.

## Experiment 1: Duration threshold measurement

### Methods

#### Participants

Six adults (2 males, 4 females, between 22-31 years) participated in this experiment.

#### Stimuli

Target stimuli had the same checkerboard patterns composed of regions with different binocular correlations as those shown in **Figure 1**. The size of the checks was 120’×120’. The complete stimulus set consisted of 4 transitions at 12 contrast levels (11.2%, 13.8%, 16.6%, 20.2%, 24.9%, 28.9%, 35.0%, 40.7%, 47.5%, 57.6%, 67.5%, 82.0%) and 7 presentation durations (16.6ms, 33.3ms, 50ms, 66.6ms, 83.3ms, 100ms, 116.6ms) for each transition type.

#### Measurement of contrast threshold for different presentation durations

Contrast threshold was measured by using an adaptive threshold search algorithm (updated maximum likelihood procedure, (19)) in a two-alternative forced choice procedure. Contrast threshold for each transition and for each presentation duration was measured in 28 separate blocks. At the beginning of the block, a magenta fixation dot indicated that the actual stimulus was being generated (i.e., idle period) and turned green when the generation was over.

Each trial within a block was composed of two intervals (**Figure 2**). The stimulus appeared for one of the 7 presentation durations in one of two equally long 2100ms stimulus presentation intervals. The first and the second stimulus intervals and the end of the trial were indicated by the change of the central fixation dot color to white, blue or green, respectively. Within each interval, the background correlogram was shown except when the target appeared or did not appear for one of the 7 predefined durations. If the target appeared, it always appeared at 1600ms from the beginning of the interval. The subject was asked to decide which interval contained the target. The contrast level for each trial in the blocks was set according to the updated maximum likelihood algorithm (19). The actual contrast threshold was defined as the inflexion point of the psychometric function fitted by the search algorithm. If the required contrast was still at maximum after 10 presentations, the block was canceled and a contrast threshold >80% was recorded. Otherwise, the contrast calculated by the search algorithm after 50 trials was taken as the final threshold. To avoid fatigue, short breaks were allowed between blocks when needed. The data set used for further analysis is shown in **S1 Table**.

**Figure 2.**
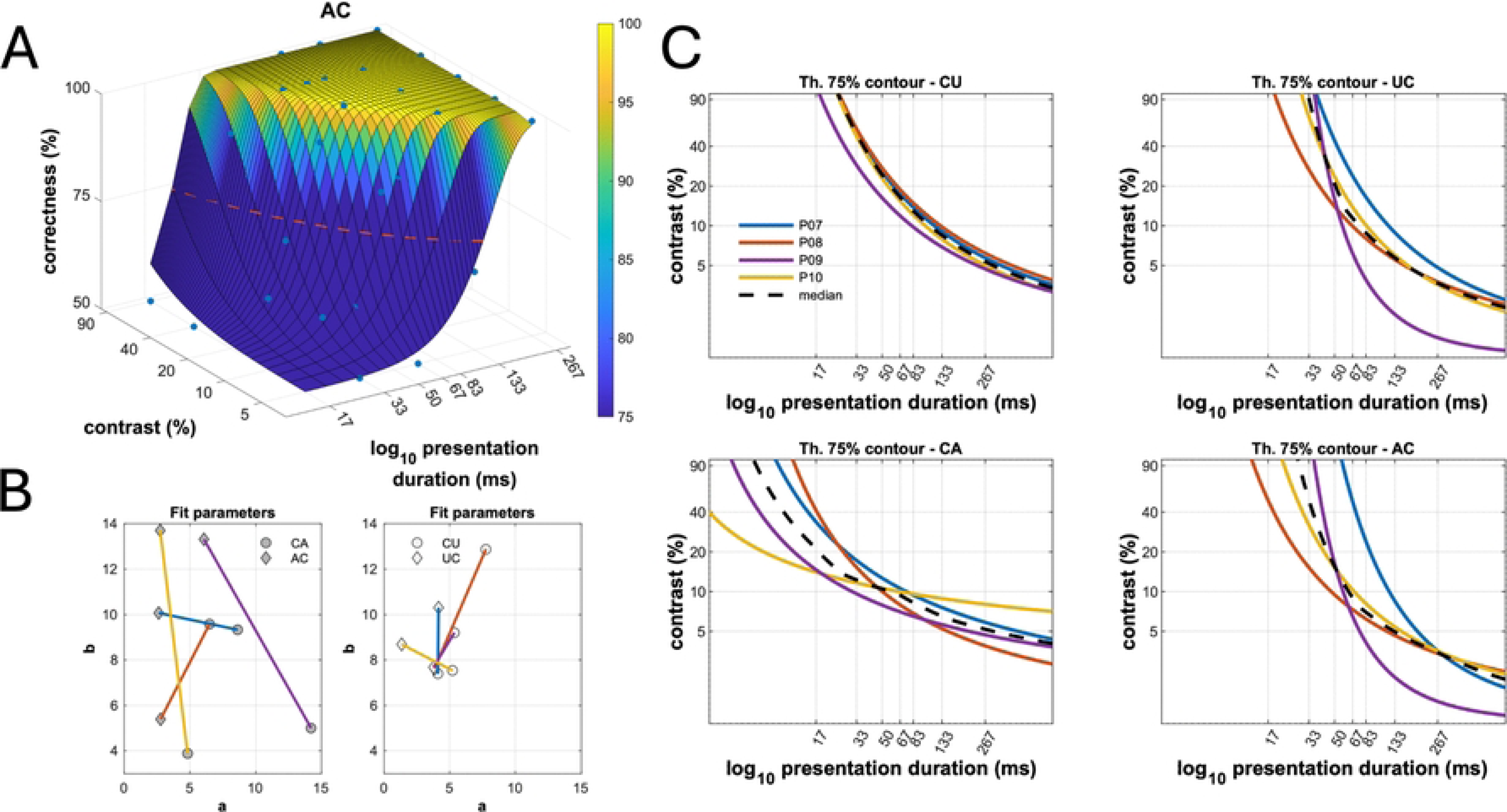
Time course of trials of the two-interval forced choice procedure for measurement of duration threshold. The example shows a CA correlation change (cf. **Figure 1**). Each trial was composed of two 2100ms long intervals that were marked for the participant by white and blue fixation dots, respectively. The stimulus appeared at 1600ms into one of two presentation intervals (top and bottom panels, respectively). The duration of the stimulus was one of the 7 presentation durations. The end of the trial was indicated by the change of the central fixation dot to green, after which the participant was asked to indicate which interval contained the target. Except for the duration of the target stimulus, the screen showed the baseline condition, i.e. in this example, a correlated DRDC. The contrast level for each trial was set according to an adaptive threshold search procedure (see text for details) for the entire trial and the preceding interstimulus interval.

#### Estimation of duration threshold of the population

Although individual duration threshold could not be determined within the above procedure, we could estimate the duration threshold characterizing all participants for each of the four transition types. For this, we determined the proportion *z* of participants that fell below a criterion contrast threshold of 25% as a function of presentation duration y (**Figure 4**). A logistic function of the following form was fitted to these data

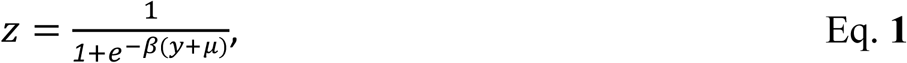

**Figure 3.**
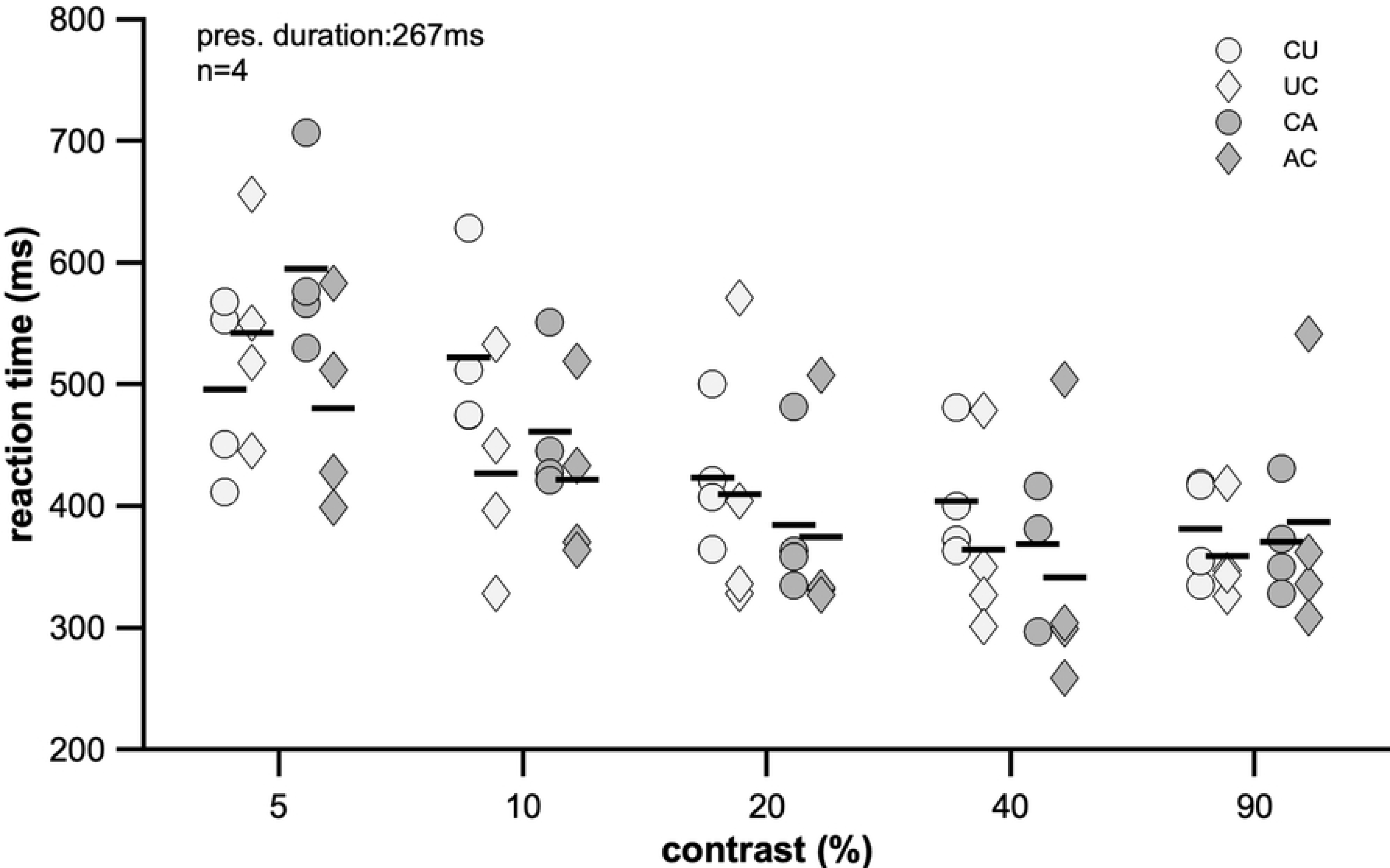
Contrast thresholds for transitions of binocular correlation presented for different durations. Each panel shows contrast thresholds as a function of presentation duration for each participant (n=6). Contrast thresholds were determined using Best PEST adaptive search algorithm. Different marker types show data for AC, UC, CU, CA transitions, respectively. Presentation duration is shown on logarithmic scale.

**Figure 4.**
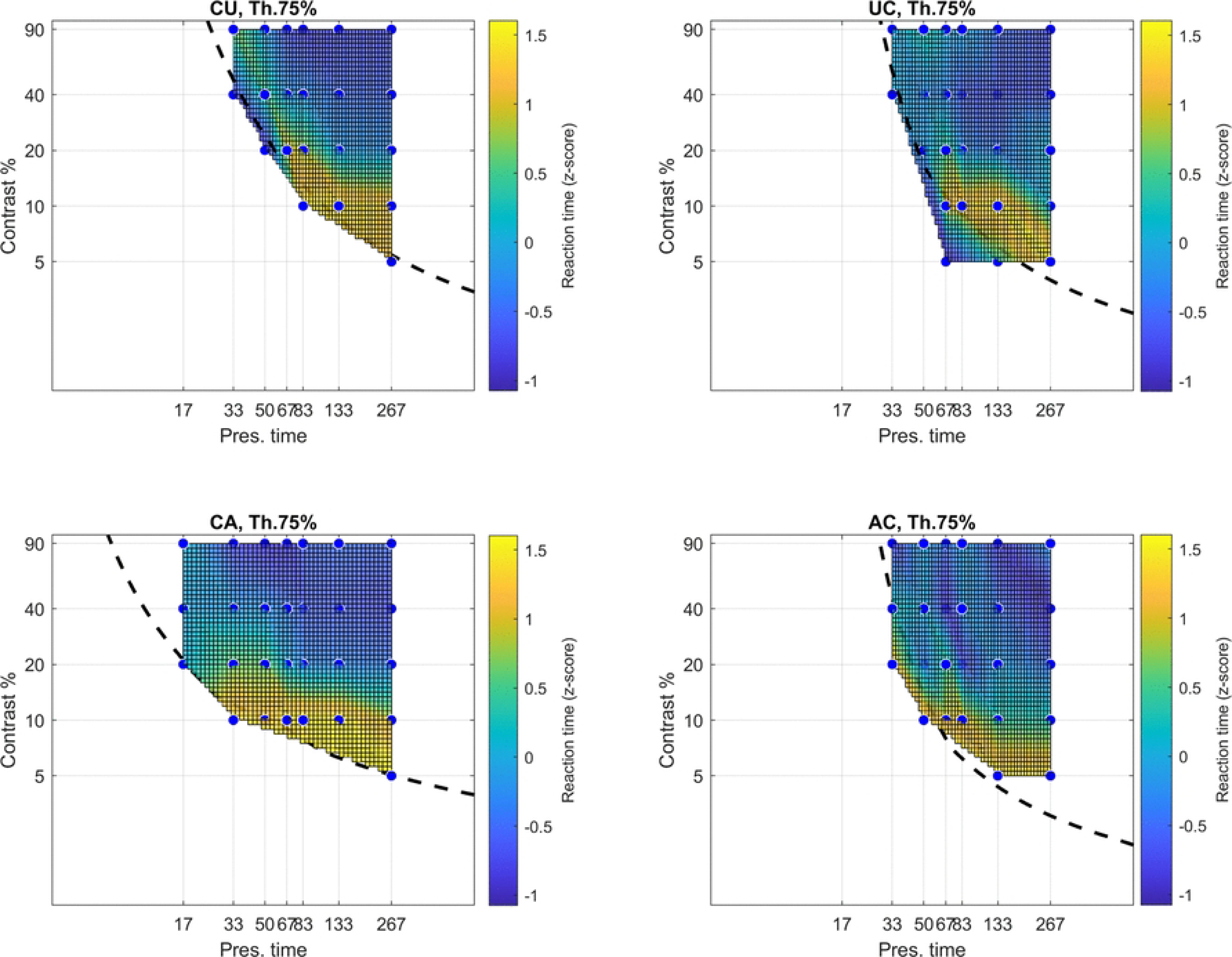
Estimating duration thresholds from Experiment 1. Percentage of participants below a criterion contrast threshold of 25% as a function of presentation duration for each of the four transition types. Lines show fitted logistic functions used to estimate the expected value and SD of presentation duration at the criterion contrast threshold.

where *μ* and *β* are the position of the inflection point and steepness of the sigmoid function, respectively. The standard deviation of the population can be determined from the steepness of the function as follows

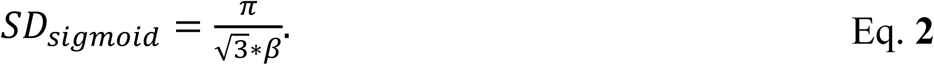

#### Results

Results are summarized in **Figure 3**–**Figure 5** (threshold values by participants; presentation duration estimation over all participants; duration threshold comparison) and **Table 1** (summary of duration thresholds). As is shown in **Figure 3**, at the shortest (i.e., 16.6ms) presentation duration contrast threshold could be measured for CA and CU transition types. However, none of the observers could detect the UC and AC transitions even at the highest available contrast level (82%). At the longest presentation duration (i.e., 116ms), all transition types could be detected at the lowest possible contrast level (11%).

**Figure 5.**
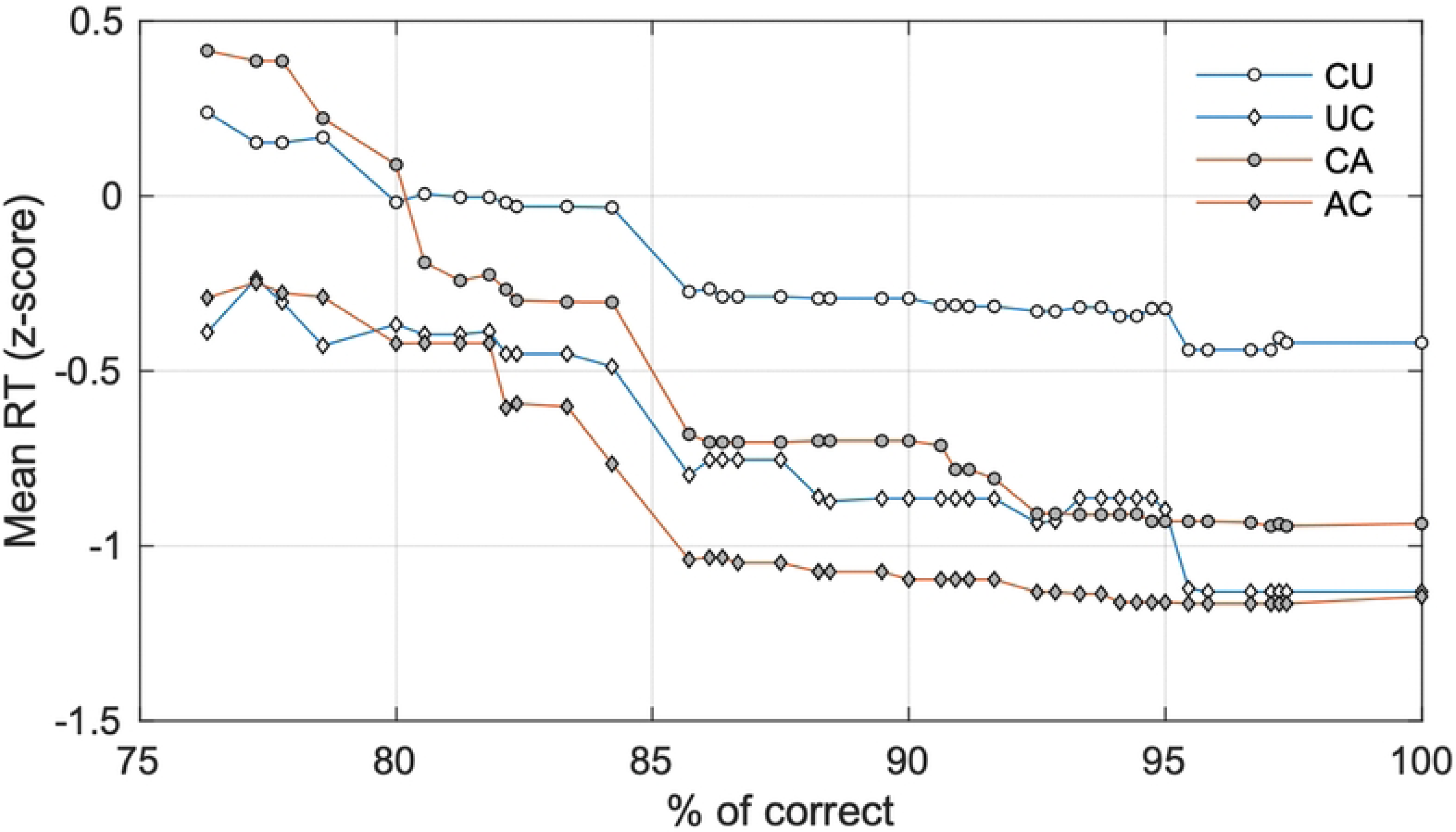
Duration thresholds from different experiments (indicated by the line type) compared. Although the three experiments were conducted in different labs and they used different paradigms, a consistent hysteresis effect is revealed.

**Table 1.**
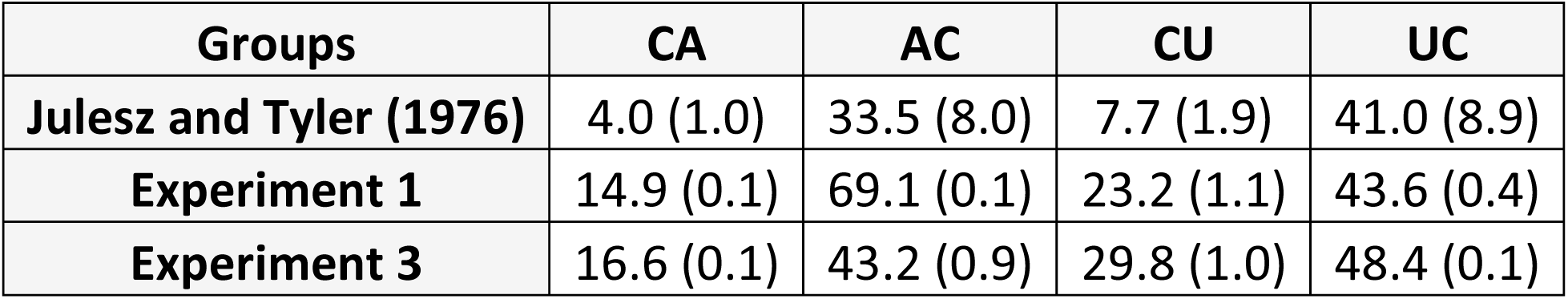
Summary of the duration thresholds (in ms) for four transition types. Mean ± SD values reported by Julesz and Tyler (11) are compared to data from the current study.

| Groups | CA | AC | CU | UC |
| --- | --- | --- | --- | --- |
| Julesz and Tyler (1976) | 4.0 (1.0) | 33.5 (8.0) | 7.7 (1.9) | 41.0 (8.9) |
| Experiment 1 | 14.9 (0.1) | 69.1 (0.1) | 23.2 (1.1) | 43.6 (0.4) |
| Experiment 3 | 16.6 (0.1) | 43.2 (0.9) | 29.8 (1.0) | 48.4 (0.1) |

According to **Table 1**, AC transition type needed the longest presentation duration (69.1ms) to be detected by the participants, and the UC transition was detected faster (43.6ms) than the AC. Both CA and CU were detected faster than their opposite counterparts, a phenomenon that we refer to as the hysteresis effect.

CA transitions were detectable within one monitor refresh cycle (i.e., 16.6ms), the shortest possible duration for our system. Since CA transitions resulted in the lowest contrast thresholds (average 20.8%) at the shortest presentation duration (16.6ms) by all subjects, the fitted psychometric function predicted the fastest (14.9ms) duration threshold. Although the CU transition was mostly also detectable at 16.6ms presentation duration, when compared to CA, it had higher contrast thresholds at both 16.6 and 33.2ms presentation duration, suggesting longer time to be recognized.

In summary, all subjects needed shorter presentation duration (PD) to detect the transition *away* from the correlated state than the opposite transitions so that duration threshold increased in the following order: CA < CU < UC < AC (**Table 1**, **Figure 4**).

Our data (**Table 1**) are not unlike those obtained by Julesz and Tyler (11) who used a similar experimental protocol although markedly different instrumentation. In order to compare our results to theirs, we calculated the mean duration thresholds of their 3 participants. The combined standard deviation (SD) of this overall mean was calculated from the SD’s of individual means using the following formula

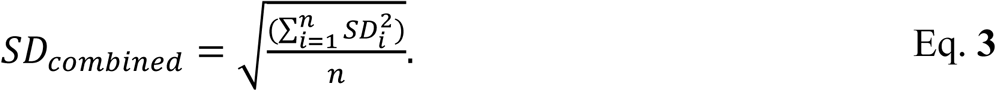

In particular, the results of Experiments 1 and 3 corroborate the presence of a hysteresis effect, a hallmark finding of the Julesz and Tyler study (7) (**Figure 5**).

## Experiment 2: Reaction time measurements

### Methods

#### Participants

Twelve adults (3 males, 9 females, 2 left-handed, 10 right-handed, between 22-31 years) participated in this experiment.

#### Measurement of reaction time

Participants were tested for the CA, AC, CU, and UC transitions at two contrast levels (90% and 10%). Each block involved two presentations of the four transitions at both contrast levels (2×4×2 conditions) in a random order. Altogether, ten reaction times (RT) were measured for each condition in 5 blocks with breaks in between when needed.

Throughout the experiment, the color of the fixation dot was changed according to the trial stage to help observers focus their attention. The trial (**Figure 6**) started with a 4s long static delay while the background was presented with a black fixation dot. When the fixation dot turned white, the observer had to pay attention to the screen expecting the target to appear. The onset of the target was jittered between 0-1s and it was displayed for 333ms in all conditions. The response key was held in the dominant hand, and the participant was asked to press the button as quickly as possible when any change in appearance - apart from the random dot refresh - was detected. The RT was measured by a custom-made microcontroller system (Arduino, Scarmagno, Italy) with millisecond accuracy, from first frame of the target until the response button was pressed. The precision of the system was tested and verified using a photodiode and an oscilloscope.

**Figure 6.**
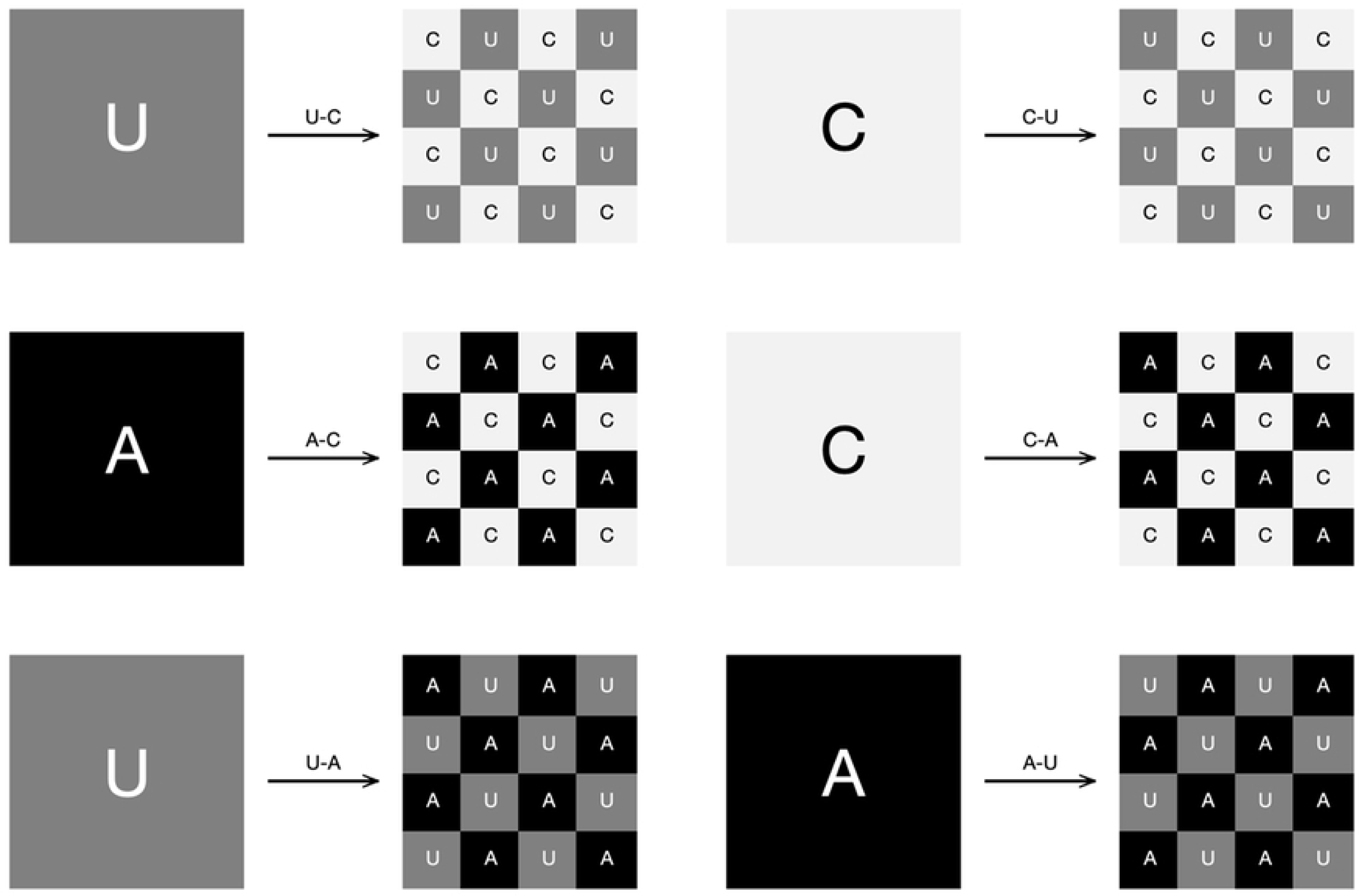
Time course of trials for measurement of simple reaction time (Experiment 2). The example shows a CA correlation change (cf. **Figure 1**). Each trial began with the central fixation dot turning from black to white. Following a random foreperiod (FP), the stimulus appeared for 333ms. Reaction time (RT) was measured from the start of the stimulus until the participant pressed response key. Except for the duration of the target stimulus, the screen showed the baseline condition, i.e. in this example, a correlated DRDC. The contrast level was either 10% or 90% for the entire trial and the preceding interstimulus interval.

RTs outside the 140-800ms range were not accepted and the same stimulus was presented again giving a second chance for perception. If an RT for a given condition was rejected twice, the “not seen” comment was recorded instead of an RT value, and the block was continued to the next stimulus. The details of the RT measurement have been published before (20). The data set used for further analysis is shown in **S2 Table**.

#### Data analysis

For each participant and stimulus condition, RT values exceeding the mean ±2 standard deviations were excluded from the statistical analysis. In some cases, Lilliefors test suggested non-normal distribution of the RTs, therefore the median RT was used characterize the response speed. To summarize RT of a given condition for all participants, the mean of the individual median RTs was used.

#### Results

The purpose of this experiment was to measure simple RTs to transitions between different types of binocularly visible correlograms (i.e., CA, AC, CU, UC) at 2 contrast levels. Repeated measures ANOVA (rANOVA) indicated a significant effect for contrast and type of change (**Figure 7**), (F(1,10)=48.833, p<.001; partial η^2^=.830; F(3,30)=10.463, p<.001; partial η^2^=.511), but not for gender (F(1,10)=1.485, p=.251; partial η^2^=.129). However, when analyzing the changes separately for the two contrast levels, a significant main effect was found only for 10% contrast ((F(3,33)=31.890, p<.001; partial η^2^=.744). At the higher contrast of 90%, RTs were statistically indistinguishable for all transitions.

**Figure 7.**
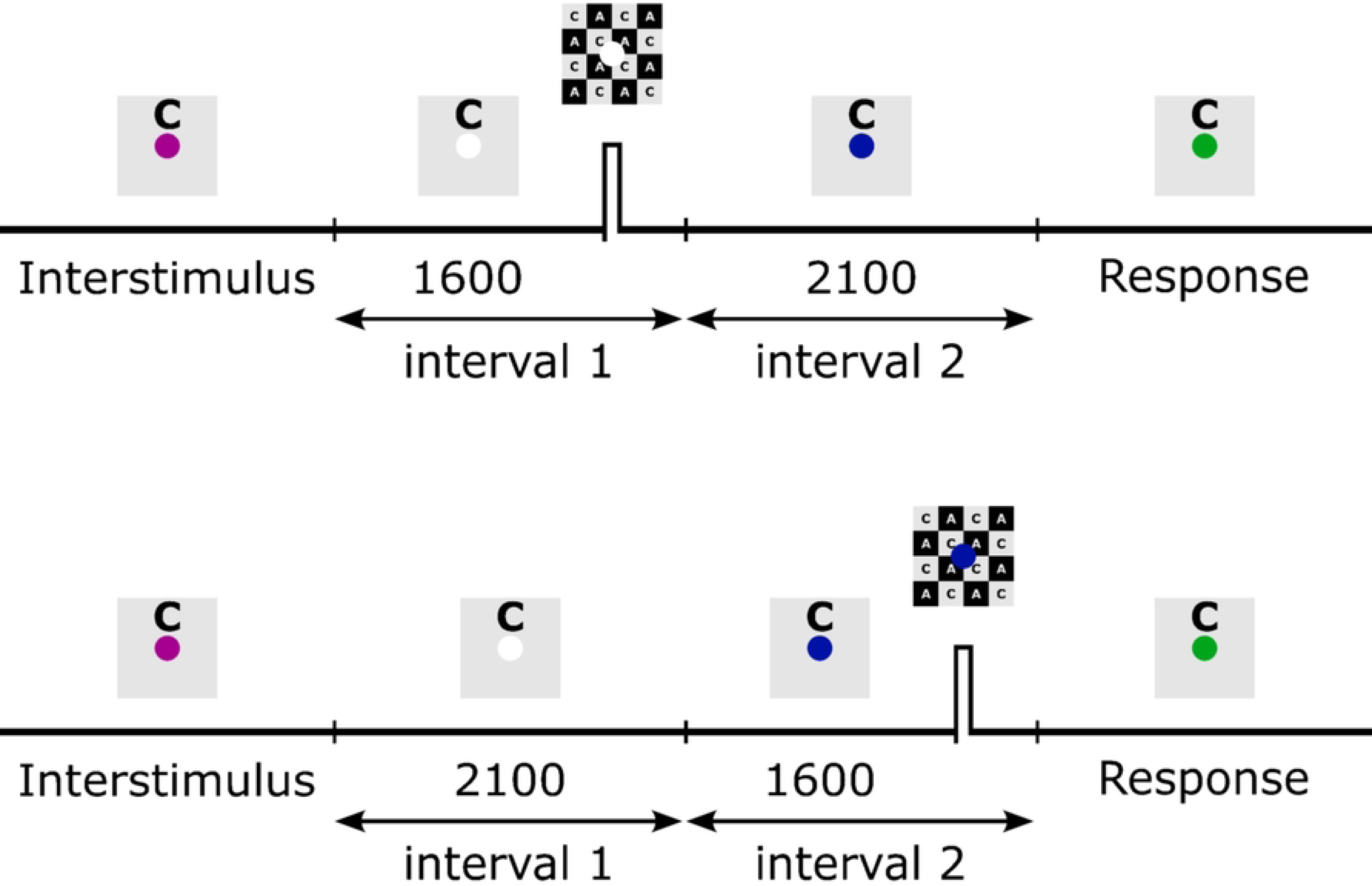
Reaction times for the different transitions at 10% and 90% contrast levels. Data points show medians of 10 presentations for each of the 12 participants. The group means are indicated by horizontal lines. Statistical comparison of these data is shown in **Table 2** and **Table 3**.

**Table 2.** Statistical comparison of reaction times at 10% contrast for Experiment 2. Reported are p-values from paired samples t-tests adjusted with Bonferroni correction. n.s., non-significant.

| 10% | Experiment 2 |  |  |  |
| --- | --- | --- | --- | --- |
|  | CU | UC | CA | AC |
| CU |  |  |  |  |
| UC | 0.009 |  |  |  |
| CA | 0.0001 | n.s. |  |  |
| AC | 0.00003 | n.s. | 0.023 |  |

**Table 3.** Statistical comparison of reaction times at 10% contrast for Experiment 3. Reported are p-values from paired samples t-tests adjusted with Bonferroni correction. n.s., non-significant.

| 10% | Experiment 3 |  |  |  |
| --- | --- | --- | --- | --- |
|  | CU | UC | CA | AC |
| CU |  |  |  |  |
| UC | n.s. |  |  |  |
| CA | n.s. | n.s. |  |  |
| AC | 0.01 | n.s. | n.s. |  |

#### Effect of contrast on reaction times

As illustrated in **Figure 7**, increasing contrast resulted in shorter RTs (p≤0.009, paired t-test), except for the AC condition (p=0.322), where no difference was found (**Table 4**). The shortening of RTs likely due to the participants approaching their asymptotic limit in RT where no stimulus related improvement was possible. Moreover, individual pairs of RTs at the two contrast levels were strongly correlated (r≥0.677, p≤0.017) suggesting contrast related differences were consistent across participants of different performance.

**Table 4.** Statistical comparison of reaction times between 90% and 10% contrast. Reported are p-values from paired samples t-tests. n.s., non-significant.

| 90% vs. 10% | Experiment 2 | Experiment 3 |
| --- | --- | --- |
| CA | 0.002 | 0.013 |
| AC | n.s. | n.s. |
| CU | $4.3 \times 10^{-7}$ | 0.024 |
| UC | 0.009 | n.s. |

#### Effect of transition type on reaction times

Due to the compression of RTs near their lowest limit at 90% contrast, the transition type did not have a significant effect (p=0.383) on RTs (**Figure 7)**. Therefore, we focus on results obtained with 10% contrast in the subsequent analysis.

The main effect for transition type at 10% was highly significant (F(3,33)=31.89, p=7.2’10-10, r=0.744). Like in Experiment 1, a significant hysteresis effect was observed: opposite transitions between the same two binocular correlation states resulted in different mean RTs. Using Bonferroni corrected pairwise comparisons, the hysteresis was significant for CA vs. AC (p=0.023) as well as for CU vs. UC (p=0.01). However, it is also evident that participants were more sensitive to transitions towards the correlated state because these resulted in shorter mean RTs than transitions from the correlated state (CA > AC and CU > UC, **Figure 7**). The difference of mean RTs was 52ms between CA and AC, and 142ms between CU and UC. Based on the pairwise comparisons of mean RTs at 10% contrast (**Table 2**), the following order of transition types can be established: AC < CA = UC < CU.

## Experiment 3: Combined detection and reaction time task

### Methods

#### Participants

Given the complexity and duration of the task, this experiment was conducted with four participants (adult females labelled P07-P10, aged between 26 and 53 years).

#### Combined measurement of contrast and duration thresholds and reaction times

Simple reaction times and duration thresholds were measured in a combined 2AFC procedure. Each target stimulus consisted of alternating stripes, each 120′ wide, composed of the baseline and target correlation states. The striped pattern was oriented either horizontally or vertically. The stimulus set contained the CA, AC, CU, UC transitions at five contrast levels (5%, 10%, 20%, 40% and 90%) and 7 presentation durations (16.7ms, 33.3ms, 50ms, 66.7ms, 83.3ms, 133.3ms and 266.7ms) at both orientations.

Each trial began with the central fixation dot turning from black to white **(Figure 8)**. Following a random foreperiod (evenly distributed between 0 and 1s), the stimulus appeared for one of the 7 presentation durations. A response key was held in the dominant hand, and the participant was asked to press the key as quickly as possible when any change in appearance – apart from the random dot refresh - was detected. Reaction time (RT) was measured from the start of the stimulus until the participant pressed the response key. Afterwards, the participants had to tell the stimulus orientation; no time limit was set for this response. Each of the 280 conditions were presented in one randomized block and each participant performed 4 to 10 blocks with breaks in between when needed. The data set used for further analysis is shown in **S3 Table**.

**Figure 8.**
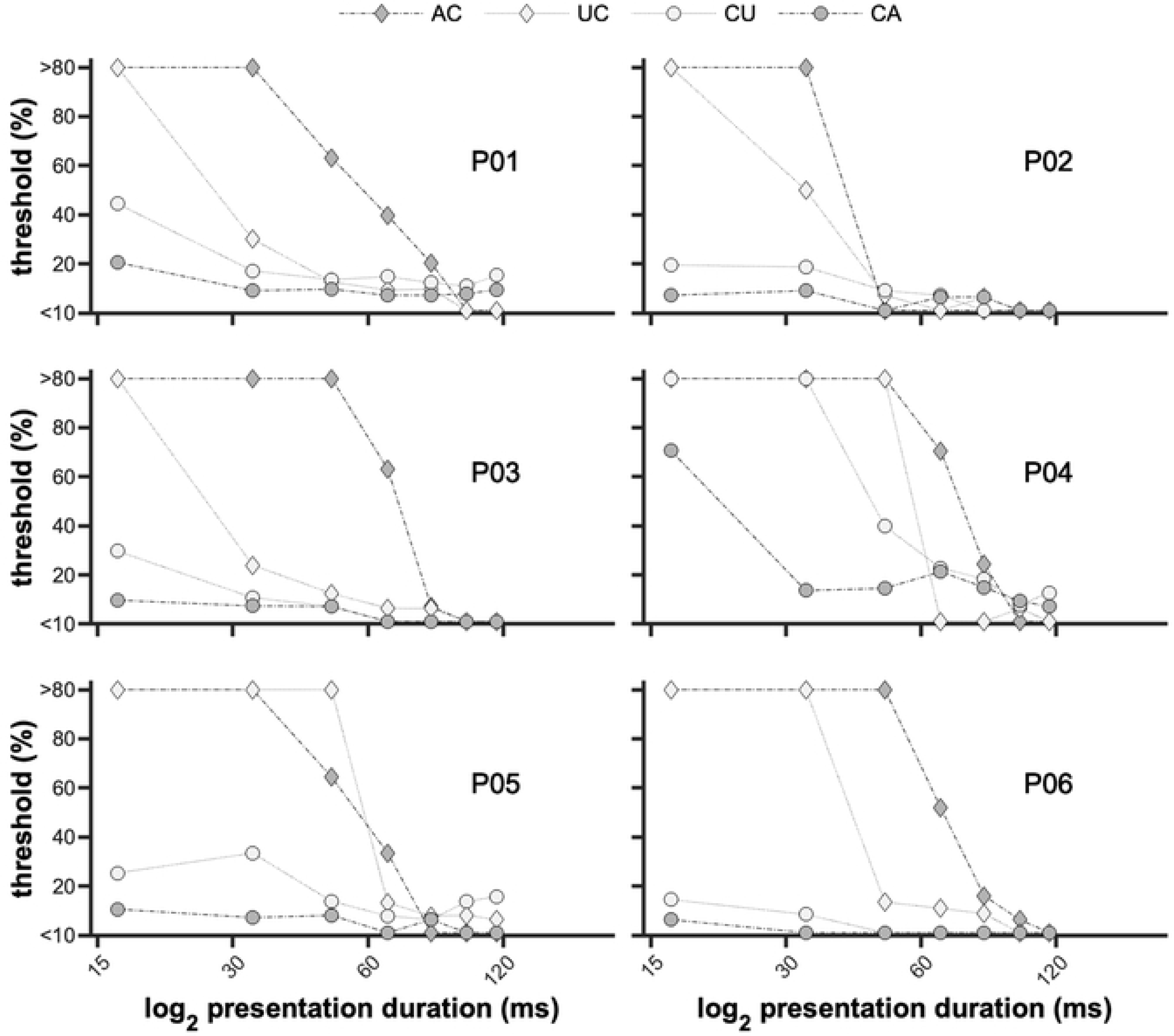
Time course of trials for combined measurement of contrast and duration thresholds and simple reaction times (Experiment 3). The example shows a CA correlation change (cf. **Figure 1)**. Each trial began with the central fixation dot turning from black to white. Following a random foreperiod (FP), the stimulus appeared for one of the 7 presentation durations. Reaction time (RT) was measured from the start of the stimulus until the participant pressed response key. The orientation of the target stimulus was either horizontal or vertical (top and bottom panels, respectively), which the participants had to indicate after pressing the response key. Except for the duration of the target stimulus, the screen showed the baseline condition, i.e. in this example, a correlated DRDC. The contrast was set to one of the 5 levels tested for the entire trial and the preceding interstimulus interval.

#### Data analysis

A bivariate psychometric function (Eq. **4**) was fitted to the proportion correct values (*z*) at each combination of presentation duration and contrast. The model function is essentially a logistic psychometric function like that in Eq. **1**, where the stimulus intensity is expressed as stimulus energy. Stimulus energy in turn, is defined as the product of (log-transformed) contrast (*x*) and presentation duration (*y*), each raised to some positive power *a* or *b*, respectively. The exponents *a* and *b* represent weighting factors by which contrast and duration contribute to stimulus energy, and *β* is a steepness factor.

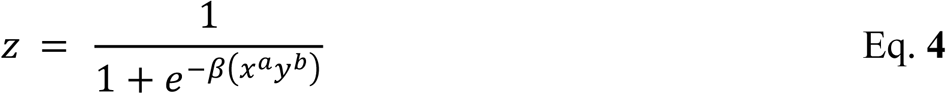

A separate psychometric function was fitted for each type of correlation change and for each participant.

Reaction times were processed and analyzed as described for Experiment 2. For **Figure 13** and **Figure 14**, each RT was expressed as z-score by calculating (𝑅𝑇 − 𝑅̅̅𝑇̅̅)/𝑆𝐷, where 𝑅̅̅𝑇̅̅ and *SD* are the mean and standard deviation of all valid reaction times for the given participant, respectively. Expressing RT as z-scores helped reduce individual differences of the mean RT between participants.

**Figure 9.**
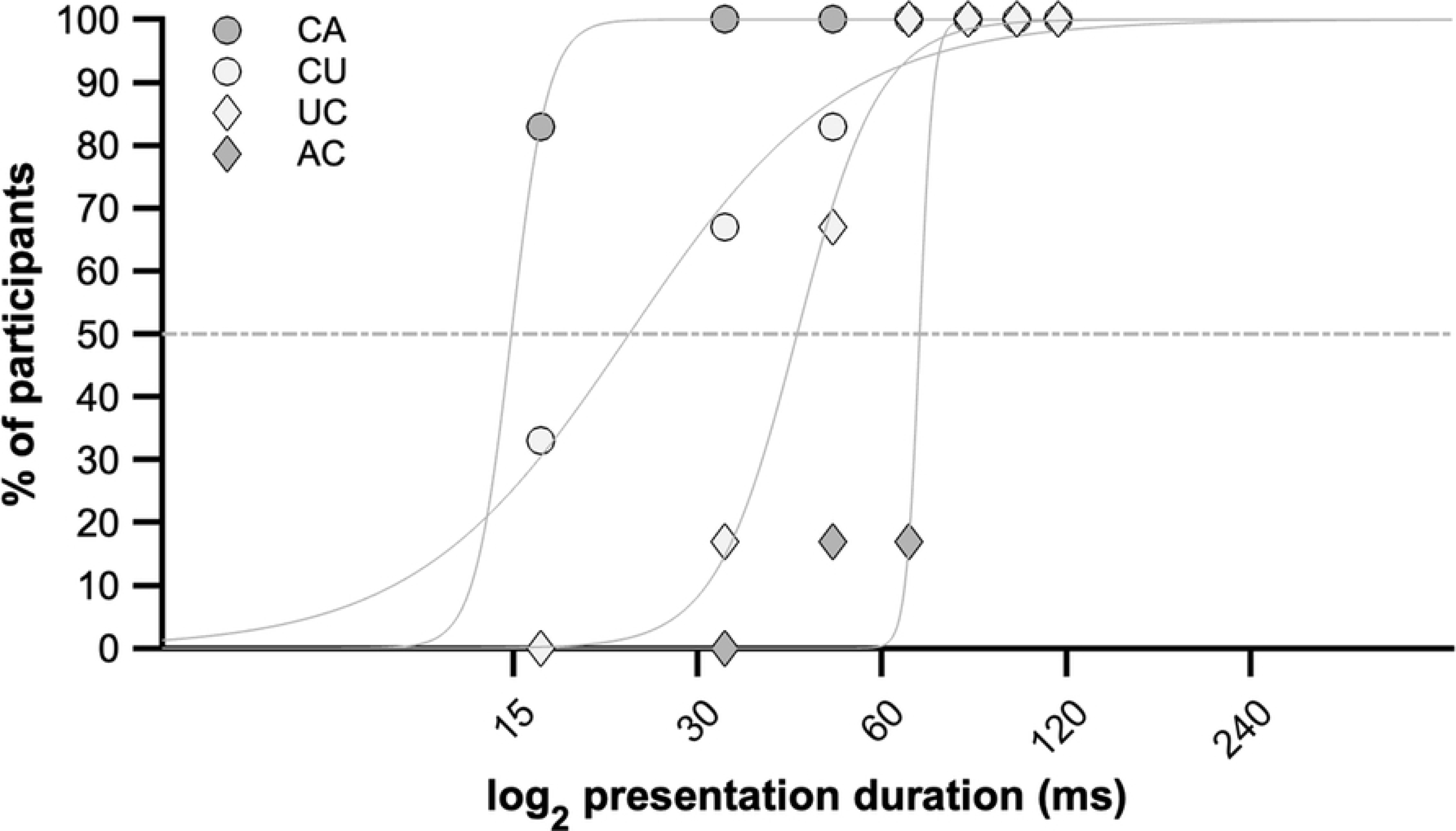
Contrast thresholds for transitions of binocular correlation presented for various durations from Experiment 3. Like in Figure 3, each panel shows contrast thresholds as a function of presentation duration for each participant (n=4). Threshold was defined as the contrast where 70% or more responses were correct. Different marker types show data for AC, UC, CU, CA transitions, respectively. Presentation duration is shown on logarithmic scale.

**Figure 10.**
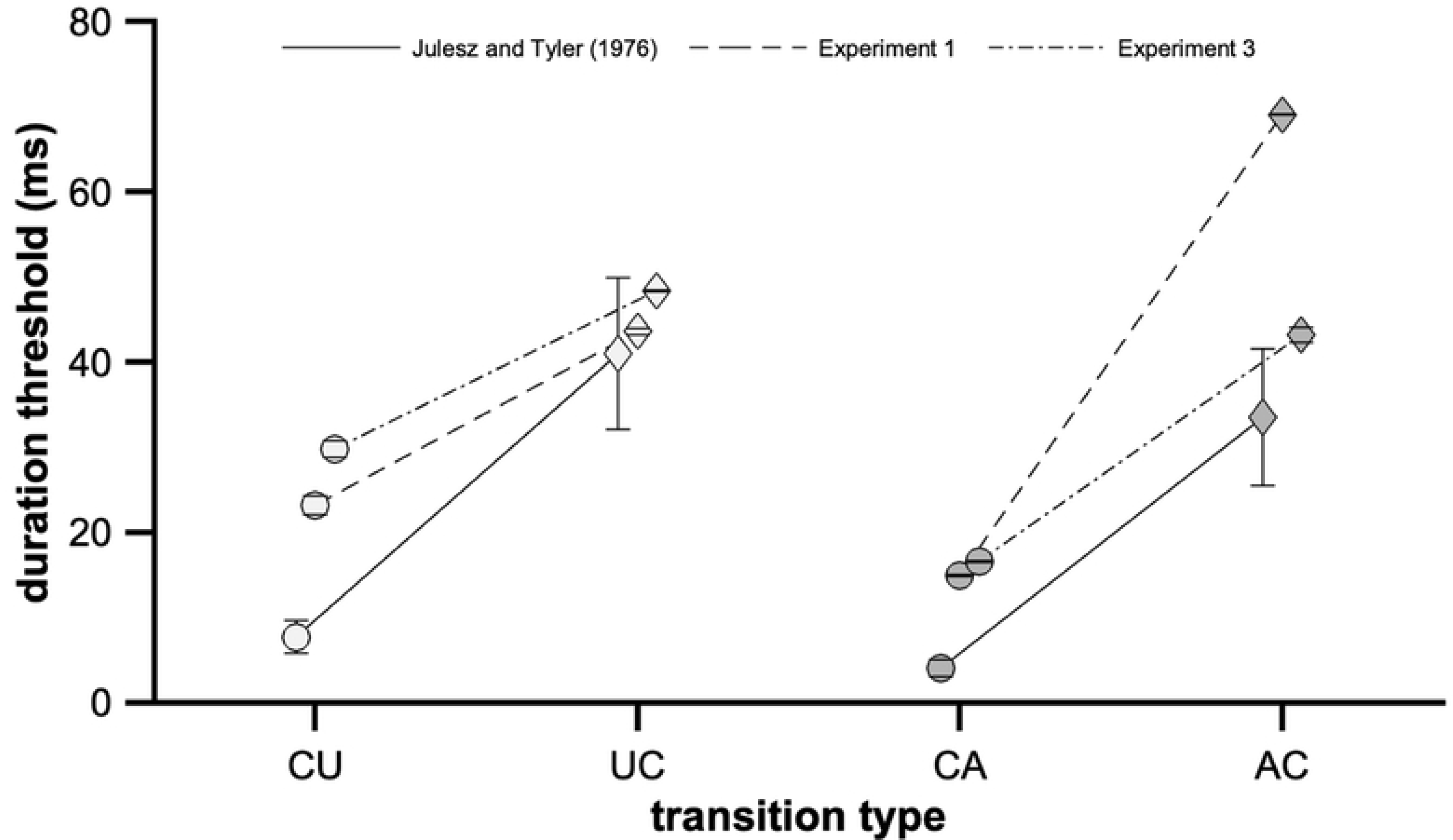
Estimating duration thresholds from Experiment 3. Percentage of participants below a criterion contrast threshold of 25% as a function of presentation duration for each of the four transition types. Lines show fitted logistic functions used to estimate the expected value and SD of presentation duration at the criterion contrast threshold like in Figure 4. Presentation duration is shown on logarithmic scale.

**Figure 11.**
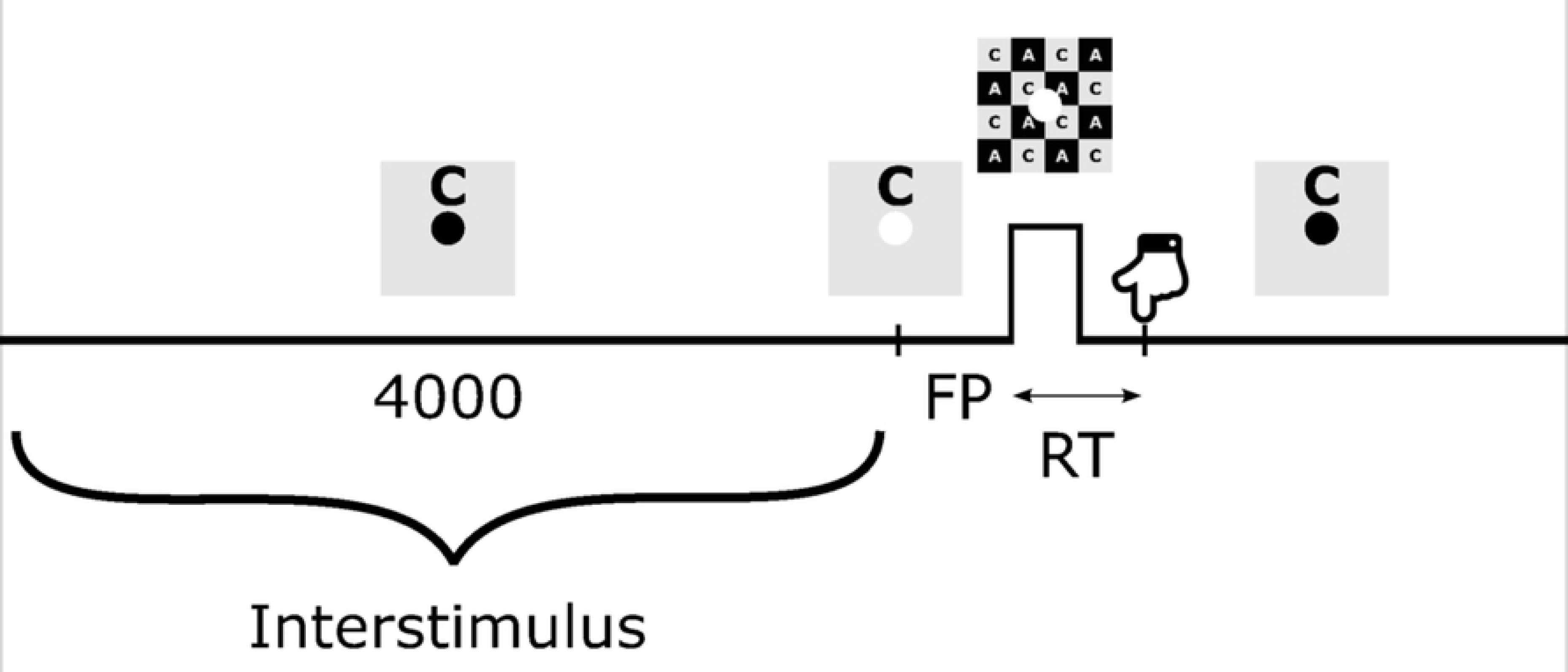
Combined measurement of contrast and duration thresholds in the 2AFC orientation detections task of Experiment 3. **A**, Proportion of correct responses as a function of presentation duration and contrast in the case of a change from an anticorrelated to a correlated DRDC for participant P09. The colored surface represents the bivariate psychometric function (Eq. **4**) fitted to the proportion of correct responses of the participant (blue dots). The estimated detection threshold for the combination of the independent variables was found as the contour at 75% correct responses (red dashed line). **B**, Comparison of the contrast and presentation duration weight parameters (*a* and *b*, respectively) of the psychometric function for each participant. The points representing parameter pairs for opposite changes (CA vs. AC and CU vs. UC) are connected to highlight their relationships in the case of each participant (color coded as in C). **C**, Threshold contours of the kind shown in **A** are plotted for combinations of presentation duration and contrast for each of the four transitions and participants (shown by different line colors). Dashed black lines denote the median threshold contours. Note the systematic right-downward shift of these contours when the change is from a baseline state of lower binocular correlation (U or A, panels on the right) compared with a change from a correlated baseline (C, panels on the left).

**Figure 12.**
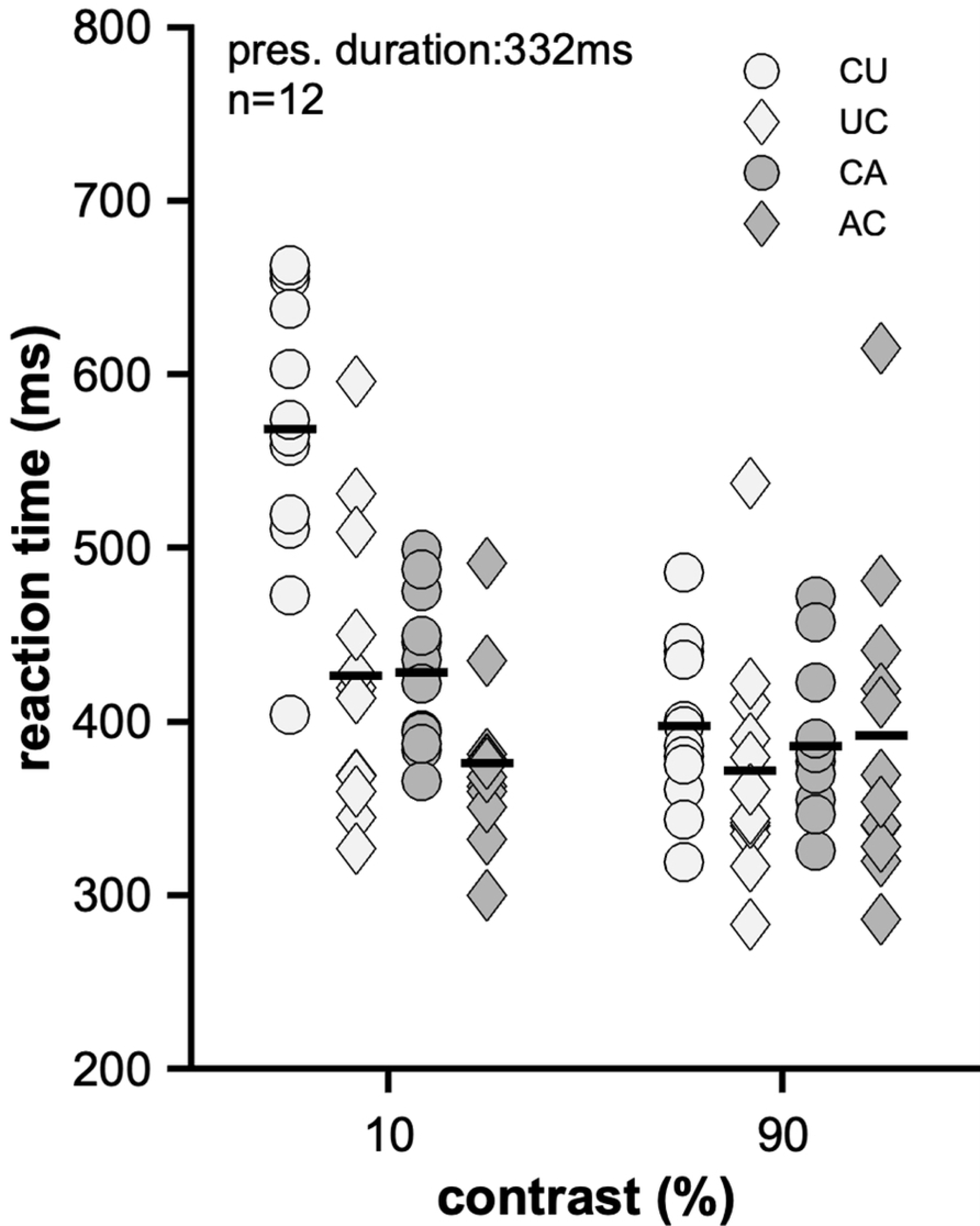
Reaction times for the different transitions at 10, 20, 40 and 90% contrast levels from Experiment 3. Data points show individual medians (n=4), the group means are indicated by horizontal lines. Statistical comparison of these data is shown in **Table 2** and **Table 3**.

**Figure 13.**
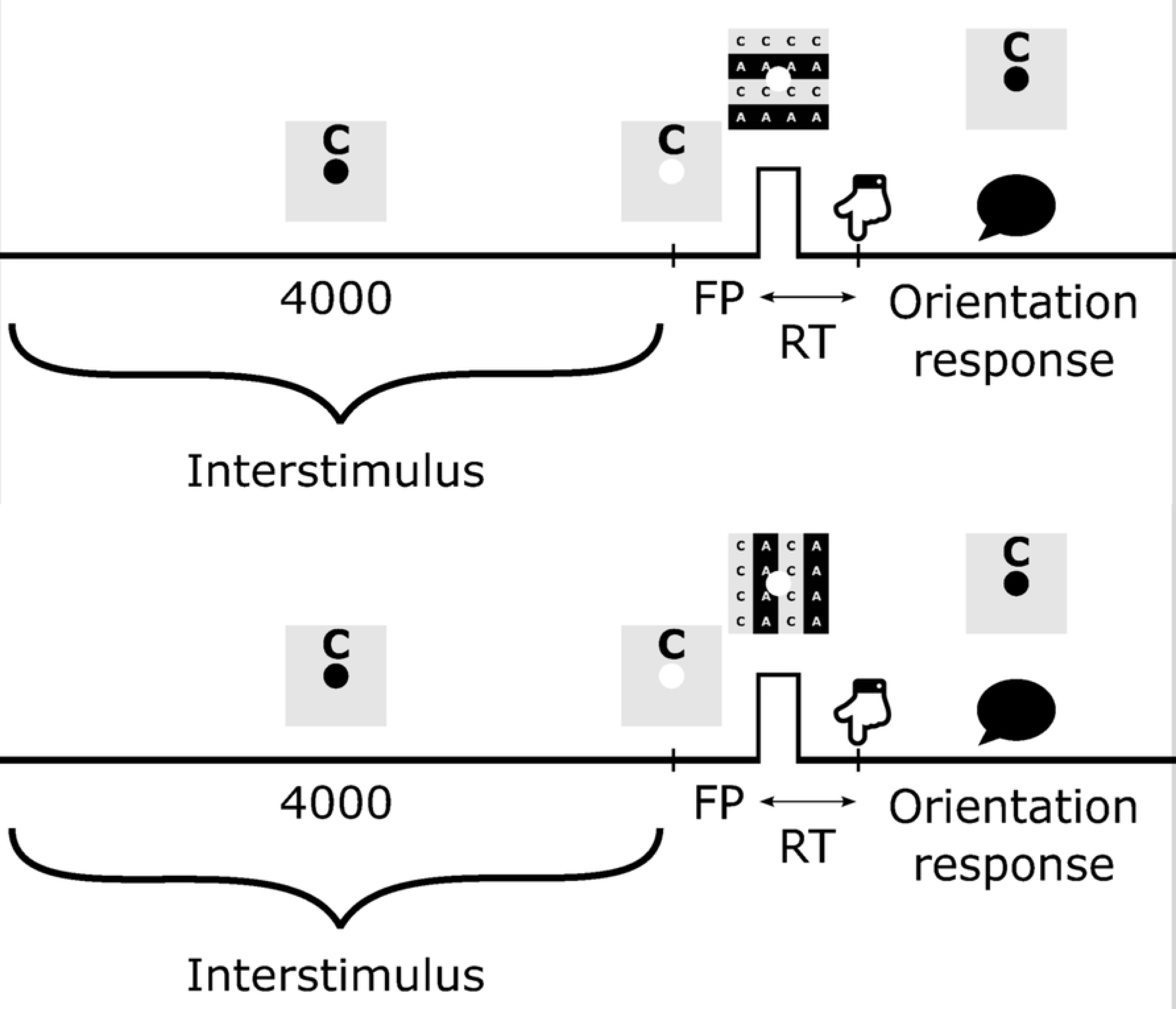
Simple reaction time z-scores as a function of presentation duration and contrast for each transition type. Color shows the mean z-score of RTs of the four participants. The dashed lines show the same median threshold contours as in Figure 11. RT z-score values are interpolated across the region where stimuli were tested, and correct detection of stimulus orientation was above threshold (>75%). The region outside is shown in white. Grid lines indicate the stimulus values tested; those combinations eliciting responses above threshold are denoted by blue dots.

**Figure 14.**
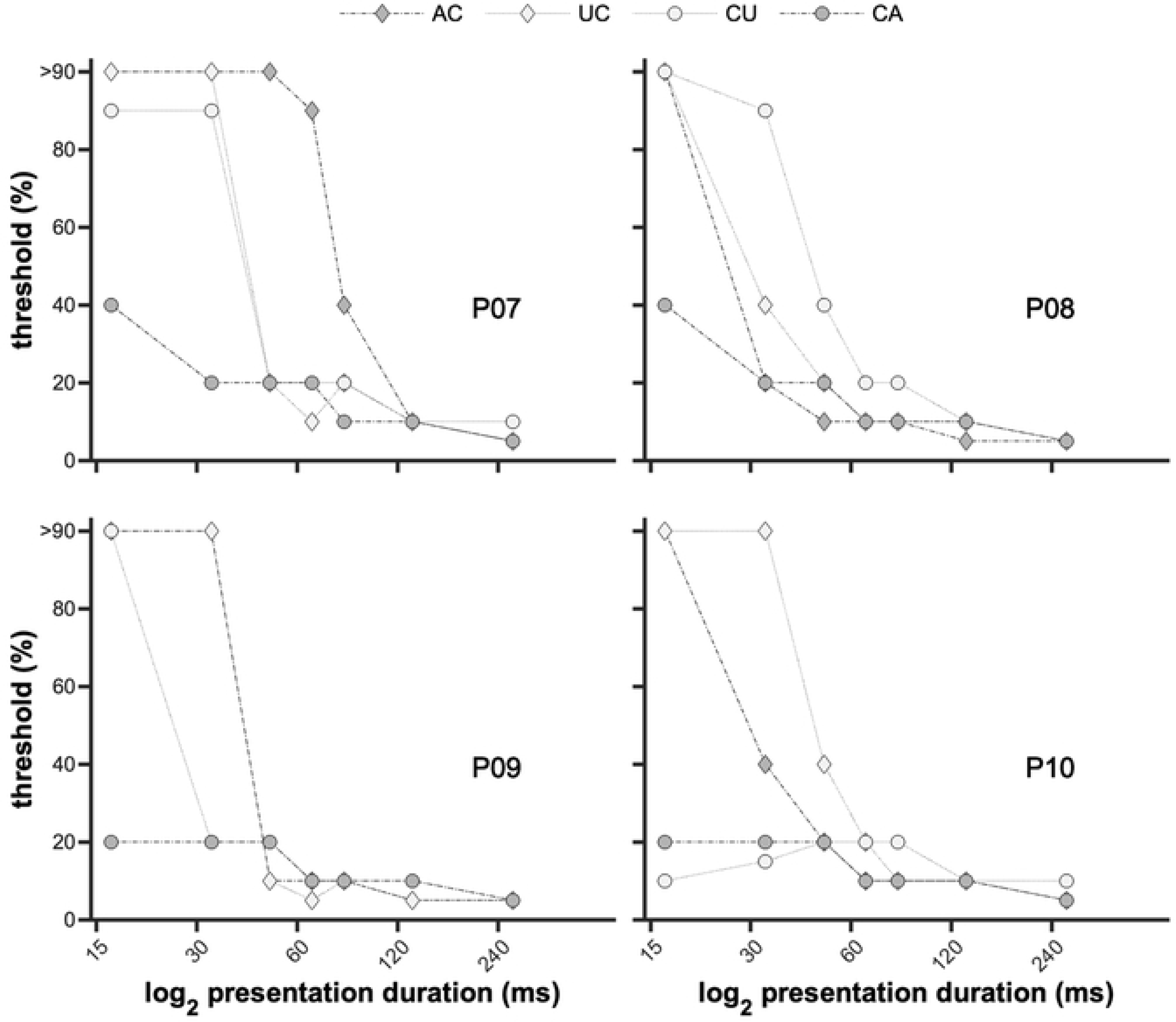
Reaction time versus percent of correct for each transition type. The curve is smoothed so that each data point with a percent correct value of x represents the mean z-scored reaction time calculated for conditions where the participants gave correct responses between x and x+15%.

### Results

#### Duration thresholds

The data from Experiment 3 were analyzed first to test if the findings from Experiment 1 can be replicated. **Figure 9** shows contrast thresholds as a function of presentation duration in a similar manner to **Figure 3**. For **Figure 9**, threshold was defined as the contrast where 70% or more responses were correct. This was done to overcome the limited resolution of the threshold estimation due to the five predefined contrast levels in this task, whereas in Experiment 1, the adaptive Best PEST procedure delivered thresholds on a finer scale. Nevertheless, the duration thresholds for each transition type could be estimated the same way as in Experiment 1 (**Figure 4**), by fitting logistic functions to the proportion of participants below a 25% criterion threshold (**Figure 10**) and using the location of the inflection points as duration thresholds. These results confirm the hysteresis effect (CA < AC and CU < UC) consistently found in Experiment 1 as well as the Julesz and Tyler (1976) (11) study (**Table 1**, **Figure 5**).

#### Detection threshold as a function of duration and contrast

So far, we analyzed detection thresholds as a function of a single variable, target duration. Experiment 3, however, allowed to examine how duration and contrast jointly determine detectability. Plotting the proportion of correct orientation judgements as a function of both variables (blue dots in **Figure 11A**) immediately reveals that contrast and duration increase detectability in a cooperative manner. This pattern is consistent with the idea of temporal integration of contrast-related stimulus intensity, where longer presentation allows the binocular mechanisms to accumulate information.

To model this joint dependence, we fitted a bivariate logistic-type psychometric function to each participant’s data for each transition type (Eq. **4**). In this framework, contrast and duration combine multiplicatively to yield an internally scaled measure of stimulus energy. This internal variable then drives the decision process through a monotonic psychometric function. An example fit is shown for participant P09 and the AC transition in **Figure 11A**.

Figure 11C shows the resulting 75%-correct threshold contours in the contrast×duration plane. The most prominent differences emerge when comparing the median threshold curves for CA and AC transitions. Both transitions exhibit a similar contrast threshold of approximately 10% at a duration of around 60 ms, where the two median contours would cross. Although the exact crossing point varies across participants, the relative relationship between CA and AC contours is highly consistent.

For durations below the crossing point (i.e., < 60 ms for the median curves), CA transitions showed lower contrast thresholds than AC transitions. Put differently, AC transitions required longer stimulus presentation to reach the same level of performance at a given contrast. This indicates reduced sensitivity to changes in correlation when the target followed an anticorrelated background stimulus.

For durations above the crossing point, this relationship reversed: CA transitions now required higher contrast to achieve threshold performance than AC transitions. This suggests that, once stimulus duration exceeds a critical integration window, the binocular system enters a more sensitive state for AC transitions than for CA transitions.

The pattern was similar although less pronounced for the CU and UC transitions. This can be explained by the smaller change in binocular correlation in these conditions, which required longer integration times to reach threshold at any given contrast. This is evident in right-shifted threshold contour of CU, when compared to its counterpart CA. In other words, when the binocular correlation change was modest, additional temporal evidence had to accumulate before the internal signal exceeded threshold, leading to the delayed (rightward) threshold curve in the contrast×duration plane. For the UC transition, increased contrast sensitivity could be observed for the longest durations: a pattern similar to that seen for AC.

#### Reaction time as a function of duration and contrast

In this section, first we analyze reaction times (RTs) in a similar manner to Experiment 2 to test if its results could be replicated under the conditions of Experiment 3. Figure 12 plots the observed median RTs for each participant, as a function of contrast and transition type. The presentation duration of 267ms was chosen for this analysis because this was closest to the value used in Experiment 2 (332ms). At this duration and 10% contrast (also used in Experiment 2), all participants could detect the orientation of the target stimulus with high confidence (100% for all transitions except for UC, where 93% of responses were correct).

Elevation of contrast to 90% resulted in saturation of RTs at their lowest asymptotic values, which did not differ significantly between transition types (repeated measures ANOVA, see General Methods). Between 10% and 90% contrast, the contrast-related decrease in mean RT was significant for CU and CA transitions, as also seen in Experiment 2 (paired t-test, **Table 4**, Figure 12). No such decrease was seen for UC and AC suggesting that the lowest possible RTs were achieved at lower contrasts for these transitions.

Lower contrast levels resulted in larger RT differences between transition types. Repeated measures ANOVAs (RMANOVA) testing for the effect of transition type in each contrast group established a significant effect at the two lowest contrast levels only; 10% (F(3,9)=15.829, p<0.001, partial η^2^=0.841) and 5% (F(3,9)=6.816, p=0.011, partial η^2^=0.694).

The relationship between the mean RTs of opposite transitions was the same as for Experiment 2 (AC < CA and UC < CU). Pair-wise statistical comparisons of the mean RTs however, showed a significant difference only for AC vs. CU (p=0.01, **Table 3**), which was nevertheless consistent with Experiment 2 (**Table 2**). This means that the hysteresis effect was not proved in the raw RTs from in this experiment, likely due to the low participant number (n=4).

In a subsequent analysis, RTs were transformed into z-scores (calculated for each participant separately) to control for individual differences better. The joint dependence of the mean RT z-scores on contrast and presentation duration is illustrated in Figure 13. For suprathreshold stimuli, RT declined with distance from the 75% threshold contour (also seen in Figure 11). As judged from the width of the yellowish region along the threshold contour, this decline was steeper for the AC transition than it was for CA. At the 10% contrast level for instance, the mean RT z-scores remained relatively high until the longest durations for CA, whereas they declined clearly for the opposite transition (Figure 13).

The complex dependence of RT on presentation duration and contrast is summarized in Figure 14. Here, RT z-scores are plotted against the proportion of correct responses. Detection probability is used here as a surrogate for internal stimulus intensity, which was in turn, a combined function of contrast and presentation duration. The relationship of the curves indicates that at any suprathreshold level of stimulus intensity, the RTs for transitions towards the correlated state (UC or AC) are shorter than RTs for the opposite transitions (CU or CA).

Thus, hysteresis of reaction times was present across all suprathreshold levels of stimulus intensity (Figure 14).

## Discussion

The effects of binocular anticorrelation might seem paradoxical because targets that transition from an anticorrelated background to a correlated pattern (AC) required a longer presentation to be perceptually detected, yet once detectable, they elicited the *fastest* reaction times (RTs) among all conditions. In contrast, transitions from a correlated background to an anticorrelated target (CA) showed the opposite pattern: they were detected with the shortest exposures but produced the slowest RTs. Overall, changes *toward* a binocularly correlated state (AC, UC) yielded significantly shorter RTs than changes away from it (CA, CU). This dissociation between detection latency and motor response speed is central to the present findings.

### Inhibition from anticorrelation and rebound disinhibition

In order to interpret our findings, it is worthwhile to consider first the interocular interactions during the background condition, from which the transitions were initiated in our experiments. Whereas the correlated state clearly set the binocular system in the state of fusion, the dynamic anticorrelated and (to a lesser extent) uncorrelated states created interocular conflict. Similar situations arise in two related experimental paradigms, namely binocular rivalry and simultaneous dichoptic masking (21–23) There is ample evidence that suppressive interocular interactions play a role in these paradigms, although their manifestation differs across conditions. In rivalry, perception oscillates between left and right eye’s inputs (24), or (under suitable conditions) the fused cyclopean (12). In cases where there is an imbalance between the two eyes’ inputs, one eye tends to dominate the percept in a temporary (e.g. flash suppression (25–27) or dichoptic masking (21–23) or permanent way, such as in amblyopia. In contrast, a dynamic anticorrelated image pair, maintains a steady state balance where neither percept can override the other, so a seemingly fused, indeed cyclopean percept arises, which Julesz described as “stereoscopic emptiness” or “woolly depth” (28). It has been long assumed that the same interocular suppression mechanisms are nevertheless active (11), although the evidence has remained indirect.

Another line of evidence, which is relevant to our study, is concerned with the perceptual phenomena around the time when there is a sudden change in binocular interactions. Classical stereoscopic adaptation studies, such as Tyler (1975) (29), demonstrated tilt and spatial frequency aftereffects following prolonged (tens of seconds) adaptation to random-dot stereogratings. Our background stimuli were viewed for less than 5 seconds before the appearance of the target, and different background conditions were interleaved within the blocks, therefore, the present findings reflect rapid, sub-second state-dependent modulation rather than long-term adaptive recalibration of disparity-selective mechanisms.

The present experiments bear more resemblance to masking paradigms. Dichoptic masking with non-cyclopean stimuli has shown strong backward masking effects, typically peaking at short stimulus onset asynchronies (∼40ms). However, such paradigms involve monocular feature interactions and are not directly comparable to the purely cyclopean transitions employed here.

Lehmkuhle & Fox (1980) (30) studied masking interactions between successive cyclopean random dot stereogram stimuli (Landolt-C target and a non-overlapping annular mask). Although background masking was also present, the prevalent interaction was forward masking extending over a relatively long time window up to ∼300ms, indicating that binocular depth signals can exhibit prolonged suppressive interactions. The results of the present study are broadly consistent with such forward suppressive dynamics between cyclopean stimuli.

Much less is known about the temporal dynamics that follow the sudden appearance or disappearance of conflicting binocular input. If binocular interactions persist transiently across stimulus transitions, the perception of the target stimulus is expected to be influenced by the preceding background state.

Our findings are consistent with the engagement of an inhibitory process that delays the formation of a fused percept while viewing an anticorrelated random-dot stimulus. Psychophysically, this is evidenced by the ∼50–70ms longer duration needed for correlated targets to reach detectability when appearing on an anticorrelated background. Two pieces of evidence from our data point to the explanation that this delay is the result of prolonged interocular inhibition or suppression triggered by the conflicting (anticorrelated) binocular signals. First, the duration thresholds for the AC transtition were considerably longer (about 2.6 to 8.3 times depending on the experiment, **Table 1**) than the thresholds for the opposite (CA) transition (hysteresis effect). Second, the CU and UC transitions, where the amplitude of correlation change was half as much as for CA or AC transitions, resulted in smaller hysteresis effect (about 1.6 to 5.3 times, **Table 1**). This can be explained by less interocular conflict and consequently less suppression, in the uncorrelated background state.

Experiments 2 and 3 aimed to probe the time window following the crossing of detection threshold by measuring reaction times. A simple assumption would suggest that once detection has occurred, the motor response would be carried out irrespective of the type of the preceding background stimulus and the type of target. Since the detection of an AC transition was delayed relative to CA, the same relationship is expected for the reaction times. Paradoxically, the opposite relationship is observed: AC transitions evoke significantly shorter RTs (Figure 7, **Table 2** and **Table 3**).

The paradox can be resolved by assuming that the removal of suppression by the interocular conflict allows the visual system to respond more rapidly, resulting in faster reaction times for the now-visible correlated target than it was the case without preceding inhibition. While the precise neural mechanism remains to be established, we propose that “disinhibition” does not only liberate the visual system from prior suppression but also invokes a rebound, effectively speeding up the accumulation of evidence toward the decision threshold that triggers the motor response. This explains how reaction time can be very short even if conscious perception was initially delayed. Thus, the paradoxical AC effect can be understood as a two-stage process: anticorrelation-induced inhibition, followed by a rebound-like facilitation when normal binocular correspondence is restored.

### Alignment with the “neurontropy model” and prior studies

The duration threshold asymmetries observed in Experiments 1 and 3 closely resemble those reported by Julesz and Tyler (1976) (11), who proposed that binocular vision involves partially distinct fusion and rivalry-related mechanisms with different temporal dynamics. Their account emphasized mutual inhibition and hysteresis between these processes, a framework that is broadly consistent with the transition-dependent asymmetries observed here.

More recent work also supports the existence of specialized binocular mechanisms for detecting interocular conflict (13,31). Kingdom et al. (2019) (6) demonstrated that sensitivity to interocular luminance differences is mediated by binocular differencing channels distinct from summation mechanisms underlying fusion. Such a pathway could provide a substrate for detecting anticorrelated input and interacting competitively with fusion-related processing. In this context, our findings are compatible with the view that correlated and anticorrelated states engage partially dissociable binocular mechanisms whose interaction produces the observed hysteresis and rebound-like facilitation.

### Neural mechanisms and implications for binocular vision

The inhibitory–disinhibitory dynamics observed here can be framed within known stages of binocular processing. In primary visual cortex, many neurons respond to binocular disparity even under contrast inversion (anticorrelation), effectively signalling potential false matches. In contrast, higher visual areas show reduced sensitivity to such anticorrelated disparities, suggesting later-stage filtering or suppression of perceptually inconsistent binocular signals.

Neurochemical evidence further suggests that anticorrelated disparity can modulate inhibitory balance in visual cortex. Using magnetic resonance spectroscopy (MRS), Matuszewski et al. (2025) (14) reported changes in GABA and glutamate concentrations during viewing of anticorrelated depth stimuli, indicating a shift in excitation–inhibition balance compared to correlated viewing. Although MRS does not provide direct evidence of synaptic inhibition, these findings are consistent with the idea that conflicting binocular input engages regulatory inhibitory mechanisms whose release may transiently enhance subsequent responsiveness.

Previous electrophysiological work has shown that adaptation to binocular anticorrelation increases cortical excitability during subsequent depth processing (32). In that study, prolonged exposure to anticorrelated random-dot stereograms resulted in enhanced steady-state disparity responses during the following test period, consistent with a shift in excitation – inhibition balance after exposure to binocular mismatch.

Our behavioral findings are compatible with this interpretation. Although the temporal structure of our paradigm differed and the duration of background stimulation was shorter, the accelerated reaction times following anticorrelated backgrounds may reflect the same underlying mechanism: exposure to interocular conflict alters inhibitory balance, such that once normal binocular correspondence is restored, the system operates in a transiently heightened state of responsiveness. Importantly, the present design does not allow us to determine how long this facilitatory state persists. Thus, our results may represent a rapid manifestation of a broader adaptation-related increase in neural excitability described previously.

More broadly, these findings suggest that inhibitory control in sensory systems may not only suppress conflicting input but also dynamically shape subsequent responsiveness, highlighting a general neural strategy for stabilizing perception under conditions of sensory ambiguity. A similar temporal pattern is sometimes observed in higher-level perceptual or cognitive decision-making, where an initial period of uncertainty or conflict is followed by a rapid and confident response once a coherent interpretation is reached, suggesting that such inhibition-release dynamics may represent a recurring motif in how the brain resolves ambiguity across processing levels.

## Acknowledgements

This research was funded by

- Thematic Excellence Program 2021 Health Sub-programme (TKP2021-EGA-16) (https://nkfih.gov.hu) from the National Research, Development and Innovation Office to GJ and PB.
- National Brain Research Program 2.0 / Nemzeti Agykutatási Program 2.0 (HU) (https://agykutatas.hu/) Nr. 2017-1.2.1-NKP-2017-00002 from the National Research, Development and Innovation Office to GJ and PB
- Hungarian Research Foundation / Országos Tudományos Kutatási Alapprogramok (HU) (http://www.otka.hu/) Nr. K108747 to PB.

## Supporting information captions

**S1 Table. Data of Experiment 1 used in the analysis.** The explanation of the variable names is as follows:

ParticipantID, unique identification number of participant.

Stimulus, type of transition between background and target states, e.g. AC is a transition from an anticorrelated background to a correlated target.

PresentationTimeMs, duration of target stimulus in milliseconds.

CalculatedThresholdContrast,contrast threshold calculated by the adaptive threshold search algorithm.

**S2 Table. Data of Experiment 2 used in the analysis.** The explanation of the variable names is as follows:

ParticipantID, unique identification number of participant.

Stimulus, type of transition between background and target states, e.g. AC is a transition from an anticorrelated background to a correlated target.

ContrastPercent, stimulus contrast in percent.

ReactionTimeMs, reaction time in milliseconds; values below 140 ms were rejected during the experiment.

Session, serial number of the session for each participant.

**S3 Table. Data of Experiment 3 used in the analysis.** The explanation of the variable names is as follows:

ParticipantID, unique identification number of participant.

Stimulus, type of transition between background and target states, e.g. AC is a transition from an anticorrelated background to a correlated target.

ContrastPercent, stimulus contrast in percent. PresentationTimeMs, duration of target stimulus in milliseconds. Orientation, either Horizontal or Vertical.

ReactionTimeMs, reaction time in milliseconds; values below 140 ms or above 800 ms were excluded from the analysis.

IsRTValid, 1 if the reaction time was >=140 ms and <=800 ms, otherwise 0. IsResponseCorrect, 1 if participant response matched the stimulus orientation, 0 if not.

